# Characterization of apicomplexan amino acid transporters (ApiATs) in the malaria parasite *Plasmodium falciparum*

**DOI:** 10.1101/2021.09.08.459553

**Authors:** Jan Stephan Wichers, Carolina van Gelder, Gwendolin Fuchs, Julia Mareike Ruge, Emma Pietsch, Josie L. Ferreira, Soraya Safavi, Heidrun von Thien, Paul-Christian Burda, Paolo Mesén-Ramirez, Tobias Spielmann, Jan Strauss, Tim-Wolf Gilberger, Anna Bachmann

## Abstract

During the symptomatic human blood phase, malaria parasites replicate within red blood cells. Parasite proliferation relies on the uptake of nutrients, such as amino acids, from the host cell and the blood plasma, requiring transport across multiple membranes. Amino acids are delivered to the parasite through the parasite surrounding vacuolar compartment by specialized nutrient-permeable channels of the erythrocyte membrane and the parasitophorous vacuole membrane (PVM). However, further transport of amino acid across the parasite plasma membrane (PPM) is currently not well characterized. In this study, we focused on a family of Apicomplexan amino acid transporters (ApiATs) that comprises five members in *Plasmodium falciparum*. First, we localized four of the *Pf*ApiATs at the PPM using endogenous GFP-tagging. Next, we applied reverse genetic approaches to probe into their essentiality during asexual replication and gametocytogenesis. Upon inducible knockdown and targeted gene disruption a reduced asexual parasite proliferation was detected for *Pf*ApiAT2 and *Pf*ApiAT4. Functional inactivation of individual *Pf*ApiATs targeted in this study had no effect on gametocyte development. Our data suggest that individual *Pf*ApiATs are partially redundant during asexual *in vitro* proliferation and fully redundant during gametocytogenesis of *P. falciparum* parasites.

**IMPORTANCE:** Malaria parasites live and multiply inside cells. To facilitate their extremely fast intracellular proliferation they hijack and transform their host cells. This also requires the active uptake of nutrients, such as amino acids, from the host cell and the surrounding environment through various membranes that are the consequence of the parasite’s intracellular lifestyle. In this manuscript we focus on a family of putative amino acid transporters termed ApiAT. We show expression and localization of four transporters in the parasite plasma membrane of *Plasmodium falciparum*-infected erythrocytes that represent one interface of the pathogen to its host cell. We probed into the impact of functional inactivation of individual transporters on parasite growth in asexual and sexual blood stages of *P. falciparum* and reveal that only two of them show a modest but significant reduction in parasite proliferation but no impact on gametocytogenesis pointing towards redundancy within this transporter family.

## INTRODUCTION

Malaria parasites replicate within human erythrocytes during the asexual blood phase, which is responsible for the symptoms of the disease. Although *Plasmodium falciparum* is able to synthesize some amino acids *de novo*^1–6^ during the intraerythrocytic development, the parasite rely heavily on amino acid acquisition from its host. Amino acids are derived either from the digestion of hemoglobin endocytosed from the infected erythrocyte^1,7^ or from the uptake of free amino acids from the blood plasma^1^. Both processes are important for efficient parasite growth^8^. The parasite is able to import all 20 naturally occurring α-amino acids from the external medium and uses them for its own protein synthesis^9–12^. Especially, the import of isoleucine^13–15^ – and for some *P. falciparum* strains also methionine^13^ – is crucial for the survival of the parasite as adult human hemoglobin lacks isoleucine. Accordingly, an increase in the permeability to a range of amino acids has been reported for erythrocytes upon *Plasmodium* infection^10,11,16,17^. This is mainly mediated via the New Permeability Pathways (NPPs) established by the parasite within the membrane of the infected erythrocyte^16,17^. While the subsequent transport across the parasitophorous vacuole membrane (PVM) is linked to nutrient-permeable channel activity^18–21^ the molecular machinery responsible for the further transport across the parasite plasma membrane (PPM) is not well defined. Neutral amino acids like isoleucine and methionine traverse the PPM more rapidly than anionic and cationic amino acids, which may be coupled to the transport of other substrates like H^+^ or Na^+^ ^16,22^. However, neither the membrane transporters nor the exact mechanism(s) by which amino acids cross the PPM have been characterized so far^8,22^.

The *P. falciparum* transportome is predicted to be encoded by 144 genes^23^, of which at least eleven are classified as putative amino acid transporters^23,24^. Five of these putative amino acid transporters belong to the Apicomplexan amino acid transporter (ApiAT)^25,26^ family. This Apicomplexan-specific family of transmembrane transporters can be subdivided into the eleven subfamilies ApiAT1–11. Some of the subfamilies are lineage-specific. For instance, ApiAT4, ApiAT8, ApiAT9 and ApiAT10 are only present in the genomes of *Plasmodium spp*.^1,25^. Others, such as the most ancient variant ApiAT2, can be found in many different Apicomplexan classes^25^. The main feature of the ApiAT family is the possession of multiple, typically twelve, transmembrane domains characteristic of solute transporters with a signature sequence between transmembrane domains 2 and 3^25^. This classifies them as members of the major facilitator superfamily (MFS)^24,27,28^. However, overall they have limited sequence similarity to other known eukaryotic or prokaryotic transporters^25^.

*P. falciparum* possesses five of the eleven ApiAT subfamilies: *Pf*ApiAT2 (*Pf*ApiAT2/MFR4: PF3D7_0914700), *Pf*ApiAT4 (MFR5: PF3D7_1129900), *Pf*ApiAT8 (NPT1: PF3D7_0104800), *Pf*ApiAT9 (MFR2: PF3D7_0104700) and *Pf*ApiAT10 (MFR3: PF3D7_0312500). Previous work in the rodent malaria species *Plasmodium berghei* (*Pb*) and the related Apicomplexan parasite *Toxoplasma gondii* (*Tg*) showed that *Pb*ApiAT8 (or *Pb*NPT1) and several other *Tg*ApiATs are localized at the PPM^29–32^ and possess amino acid transport activity^25,26,29–34^. To date, only *Pb*ApiAT4 has been shown to play an important role in parasite proliferation within erythrocytes^35^. Apart from this, *Pb*ApiAT8 is essential for gametocyte development^29,30,35^ and all variants except *Pb*ApiAT2 appear to be necessary for exflagellation of male gametocytes^35^. Additionally, sporozoite development likely requires *Pb*ApiAT2, *Pb*ApiAT4, *Pb*ApiAT9 and *Pb*ApiAT10^35^ and a lack of *Pb*ApiAT2 further impairs oocyst development. Furthermore, an association of episomally overexpressed *Pf*ApiAT10-GFP (termed MFR3-GFP) with the mitochondrion has been reported^36^.

In this study, we characterize the localization and essentiality of the ApiAT family members of *P. falciparum* during intraerythrocytic development in order to further dissect the function of these unique transporters.

## RESULTS

### *P. falciparum* ApiATs localize at the parasite plasma membrane

The five members of the ApiAT family in *P. falciparum* show different gene expression patterns and mRNA levels during the intraerythrocytic developmental cycle (IDC)^37^. While *Pf*ApiAT2 has its maximum transcript level in early ring stage parasites (8 hpi) and *Pf*ApiAT4 and *Pf*ApiAT10 mRNA levels peak in late ring stage parasites (16 hpi), *Pf*ApiAT8 shows a maximum of transcripts in late stage schizonts (48 hpi) and mRNA of *Pf*ApiAT9 is almost absent during the IDC (Figure 1A). Overall, *Pf*ApiAT2 and *Pf*ApiAT4 are most abundantly expressed on mRNA level.

**Figure 1:**
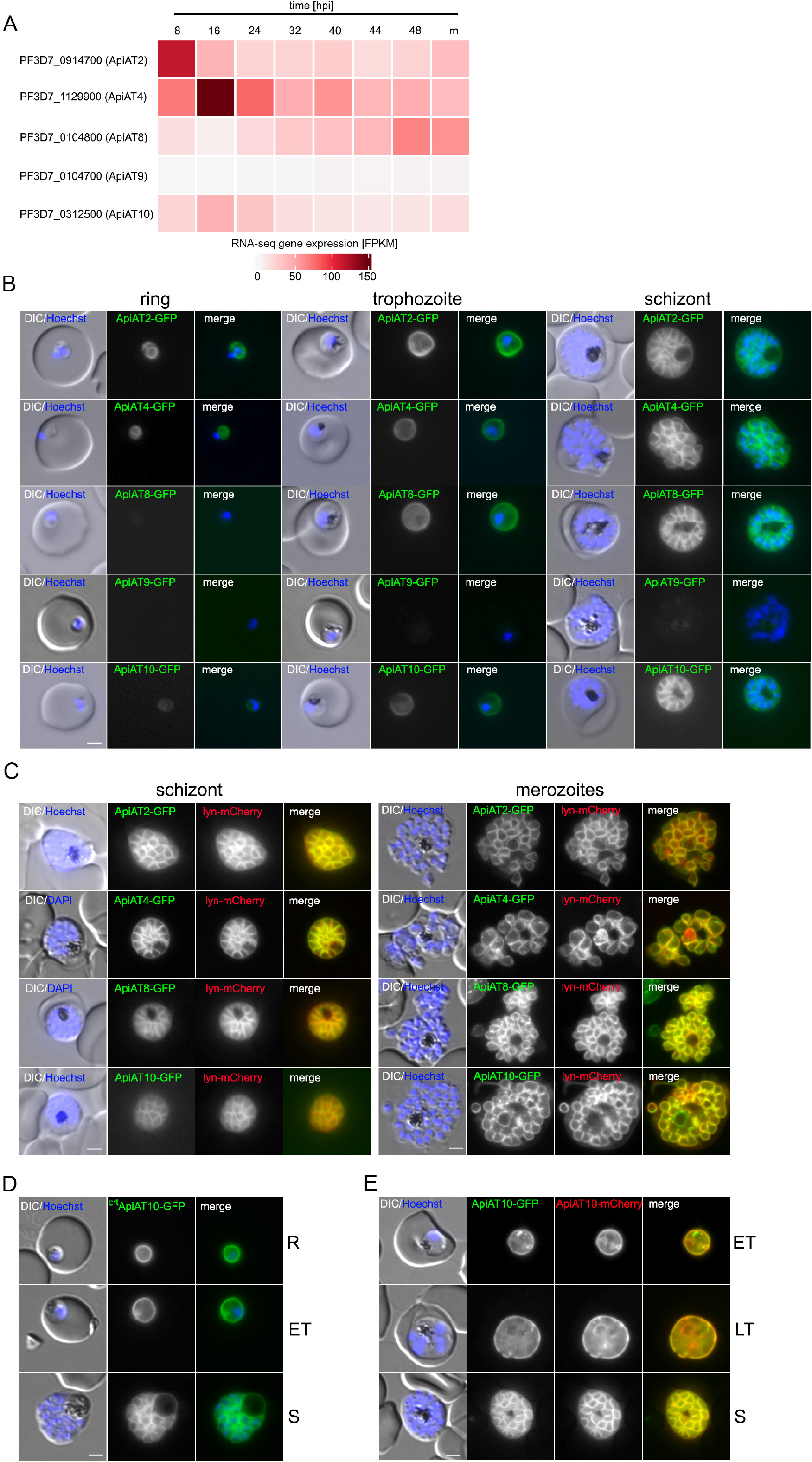
*P. falciparum* ApiATs localize to the parasite plasma membrane (PPM) during asexual blood stage development. **(A)** Heatmap of RNA-seq gene expression profiles^37^ of *Pf*ApiAT2, *Pf*ApiAT4, *Pf*ApiAT8, *Pf*ApiAT9 and *Pf*ApiAT10 during the asexual blood stage development. Timepoints indicated as hours post infection (hpi) plus merozoites (m). **(B)** Localization of *Pf*ApiAT2-GFP, *Pf*ApiAT4-GFP, *Pf*ApiAT8-GFP, *Pf*ApiAT9-GFP and *Pf*ApiAT10-GFP by live-cell microscopy across the IDC of 3D7 parasites. **(C)** Co-localization of the GFP-tagged ApiAT fusion proteins with the PPM marker protein lyn-mCherry in schizonts and merozoites. **(D)** Live-cell microscopy of 3D7-crt-ApiAT10-GFP parasites across the IDC. **(E)** 3D7-iGP-ApiAT10-GFP/ama1-ApiAT10-mCherry parasites at trophozoite and schizont stage. Nuclei were stained with Hoechst-33342. Scale bar, 2 µm. Parasite stages as indicated; ring stage (R), early trophozoite (ET), late trophozoite (LT) and schizont (S).

To determine their protein expression and localization, we tagged each of the five members of the *Pf*ApiAT family endogenously with GFP using the selection-linked integration (SLI) system^38^. Correct integration of the corresponding targeting plasmids into the respective genomic loci was verified by PCR (Figure S1A). Except for 3D7-ApiAT9-GFP, all generated transgenic cell lines expressed the full-length fusion protein (Figure S1B) to a sufficient level that allowed its subcellular localization. All of them are localized at the PPM (Figure 1B). This localization was confirmed by co-localization with the episomally expressed PPM marker Lyn^39^-mCherry^38^ and becomes particularly evident in schizont stage parasites and free merozoites, when PPM and PVM can clearly be separated (Figure 1C).

In contrast to published data^36^, we also found endogenously GFP-tagged *Pf*ApiAT10 localizing at the PPM. To re-probe into the apparent PPM localization of endogenously GFP-tagged ApiAT10 we also over-expressed this gene as a GFP and mCherry fusion protein using two different promotors (*crt*^40^ *or ama1*^41^). This allowed us to assess the influence of the tags as well as differential expression profiles on ApiAT10 protein localization. All cell lines showed PPM localization of the ApiAT10 fusion proteins (Figure 1B-E, S1). Of note, *Pf*ApiAT10-TGD parasites showed also no conferral of drug resistance to Atovaquone (Figure S1B).

### Individual *Pf*ApiATs are not essential for asexual blood stage development

To probe into the essentiality of the ApiAT family for asexual parasite proliferation, we first targeted the most abundantly expressed *Pf*ApiATs, *Pf*ApiAT2 and *Pf*ApiAT4 by conditional knockdown. Down-regulation was achieved by introducing a glmS ribozyme sequence^40^ before the 3’UTR of either the *apiat2* or *apiat4* genomic locus (Figure S1). The ribozyme was activated by addition of 2.5 mM D-(+)-glucosamine hydrochloride (GLCN), which resulted in degradation of mRNA and therefore decreased protein levels. This was assessed and quantified by live cell fluorescence microscopy after two cycles. GLCN treatment resulted in a decreased GFP-fluorescence of 85.9 % ± SD 0.9 for 3D7-*Pf*ApiAT2-GFP or 74.8 % ± SD 14.0 for 3D7-*Pf*ApiAT4-GFP (Figure 2A–D) and led to a moderate, but significant reduction of parasite growth of 30–35 % compared to the control (Figure 2E). This data indicates that *Pf*ApiAT2 and *Pf*ApiAT4 play a role in efficient blood cell proliferation but imply that they might be non-essential. Therefore, we targeted these genes with deletion constructs using the SLI system^38^ that lead to the expression of severely truncated versions of the ApiATs. In this targeted gene disruption (TGD) approach, we also included *Pf*ApiAT8 and *Pf*ApiAT10.

**Figure 2:**
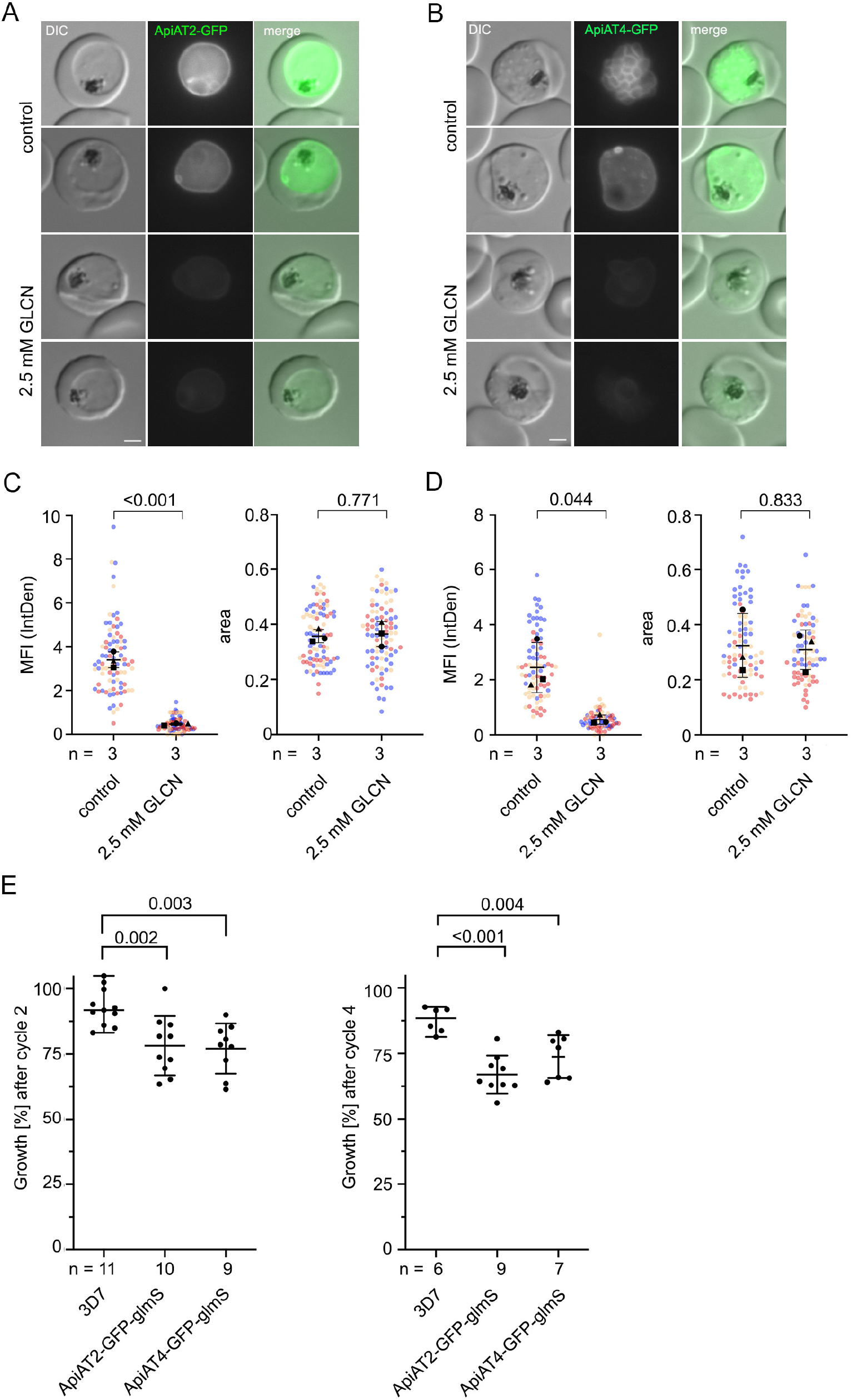
Conditional knockdown of *Pf*ApiAT2 and *Pf*ApiAT4 reveals a minor growth defect during asexual blood stage development. **(A, B)** Live-cell microscopy of **(A)** 3D7-ApiAT2-GFP and **(B)** 3D7-ApiAT4-GFP parasites, which were treated for 40 hours with 2.5 mM glucosamine (GLCN) or that were left untreated (control). Scale bar, 2 µm. **(C, D)** Quantification of knockdown efficiency by measuring mean fluorescence intensity (MFI) density and parasite size (area) of **(C)** 3D7-ApiAT2-GFP and **(D)** 3D7-ApiAT4-GFP parasites 40 hours after treatment with or without 2.5 mM glucosamine. Data is displayed as mean +/- SD of three independent experiments and individual data points are displayed on a scatterplot color-coded by experiments according to Superplot guidelines^81^. P-values displayed were determined using a two-tailed unpaired t-test with Welch’s correction. **(E)** Growth of parasites treated with or without 2.5 mM GLCN after two and four parasite replication cycles as determined by flow cytometry. Shown are relative parasitemia values, which were obtained by dividing the parasitemia of glucosamine-treated cultures by the parasitemia of the corresponding untreated ones. Displayed are means +/- SD of independent growth experiments with the number of experiments (n) indicated. Adjusted p-values displayed were determined with a two-tailed unpaired t-test with Welch’s correction and using the Benjamini-Hochberg correction afterwards accounting for multiple testing by comparing ApiAT2-GFP-glmS or ApiAT4-GFP-glmS cultured with 2.5mM GLCN to 3D7 parasites cultured with 2.5mM GLCN.

Imaging of the TGD cell lines revealed a more diffuse, but still membrane-associated GFP signal (Figure 3A–D). This might be due to the remaining transmembrane domains of the truncated *Pf*ApiAT mutants; however, our approach deleted at least ¾ of their predicted TM domains and thus most likely abolished transporter activity. In concordance with the inducible knockdown data, functional inactivation by truncation of *Pf*ApiAT2 and *Pf*ApiAT4 in the corresponding transgenic cell lines (3D7-ApiAT2-TGD, 3D7-ApiAT4-TGD) led to a moderate decrease of parasite proliferation of 20.2 % ± SD 3.2 and 19.8 % ± SD 8.6 after two parasite replication cycles (Figure 3E). No significant reduction of growth was observed upon disruption of *Pf*ApiAT8 and *Pf*ApiAT10 (Figure 3E). Interestingly, cultivation in amino acid depleted medium (approximately 90 % reduced concentration) did not indicate a higher susceptibility of any of the TGD cell lines to low amino acid concentrations compared to wild type parasites (Figure 3F).

**Figure 3:**
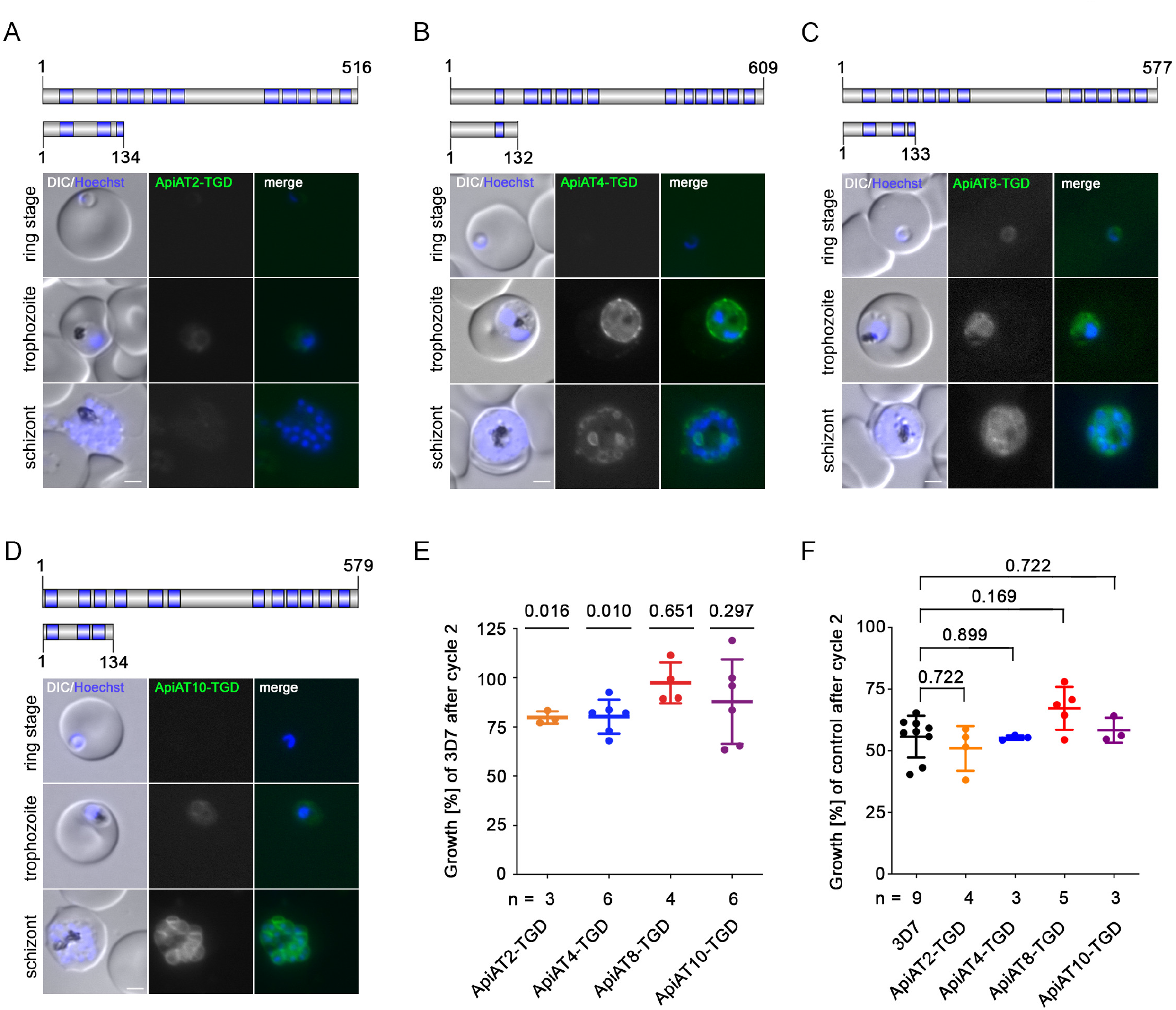
Targeted gene disruption (TGD) of *Pf*ApiAT2, *Pf*ApiAT4, *Pf*ApiAT8 and *Pf*ApiAT10 reveals the dispensability of *Pf*ApiATs for *in vitro* parasite proliferation. **(A–D)** Schematic representation of the full-length and truncated protein versions (upper panel). Protein length (number of amino acids) and putative transmembrane protein domains (blue) are indicated. Localization of **(A)** *Pf*ApiAT8-TGD-GFP, **(B)** *Pf*ApiAT4-TGD-GFP, **(C)** *Pf*ApiAT2-TGD-GFP and **(D)** *Pf*ApiAT10-TGD in ring, trophozoite and schizont stage parasites is shown in lower panels. Nuclei were stained with Hoechst-33342. Scale bar, 2 µm. **(E)** Growth of 3D7-ApiAT-TGD cell lines as percentage of 3D7 parasites growth, monitored over two intracellular development cycles by flow cytometry. The number of independent growth experiments (n) per 3D7-ApiAT-TGD cell line is indicated. 3D7 wildtype parasites were measured in parallel. Statistical differences were analyzed using a one-sample t-test with Benjamini-Hochberg correction accounting for multiple comparisons. **(F)** Growth of TGD and 3D7 cell lines cultivated in low amino acid medium relative to their growth in standard medium is shown as percentage of growth after two parasite replication cycles. The number of individual growth experiments (n) performed is indicated for each 3D7-ApiAT-TGD line, additionally 3D7 wildtype parasites were analyzed with n=9. No statistical differences were observed by comparing relative growth of TGD cell lines to 3D7 using a two-tailed unpaired t-test with Bonferroni correction.

To probe into potential transcriptional perturbations within this gene family due to functional inactivation of a single member, quantitative real-time PCR (qPCR) analysis was performed using RNA from four different time points during asexual blood stage replication. However, no consistent upregulation of RNA levels of other *Pf*ApiAT family members was observed in individual *Pf*ApiAT TGDs (Figure S3).

### *Pf*ApiATs are dispensable during gametocyte development

Previous data^30,35^ indicated a role of ApiAT8 during gametocyte development of the rodent malaria parasite *P. berghei*. Therefore, we re-engineered the GFP-tagged gene knockdown (Figure S4) and deletion cell lines (Figure S5) for *Pf*ApiAT2, *Pf*ApiAT4, *Pf*ApiAT8 and *Pf*ApiAT10 in an inducible gametocyte producing parasite line (3D7-iGP-GDV1GFP-DD^41^) using the same SLI approach. The resulting parasite lines allowed a robust, efficient and synchronized induction of gametocytogenesis and therefore a solid basis for phenotypic analysis. First, using the C-terminal GFP tag, we confirmed expression of all four *Pf*ApiATs in gametocytes. As expected, most *Pf*ApiATs remain PPM localized during gametocytogenesis, which was additionally confirmed by the colocalization with the episomally expressed PPM marker Lyn^39^-mCherry^38^ (Figure 4A–F). The exception was *Pf*ApiAT9, which showed only a faint background staining in all gametocyte stages in the 3D7-ApiAT9-GFP line (Figure 4D). The prominent GFP signal at the food vacuole in gametocytes is most likely an unspecific hemozoin signal, as it is also observed in 3D7-iGP control parasites (Figure S4C). Of note, *Pf*ApiAT2 was observed to be strongest expressed in early-stage gametocytes and weaker in late-stages (Figure 5B, D; Figure S4D), while *Pf*ApiAT4 showed strongest expression in late-stage gametocytes (Figure 5C, E). Next, we investigated the consequence of glmS-based conditional knockdown for *Pf*ApiAT2 and *Pf*ApiAT4 (Figure 5A). Although 75–80 % knockdown of *Pf*ApiAT2 or *Pf*ApiAT4 expression was achieved (Figure 5B–E), no significant reduction in gametocytemia or aberrant gametocyte development could be detected (Figure 5F). This was re-investigated by targeted gene disruption. Likewise, deletion of these two genes as well as of *Pf*ApiAT8 and *Pf*ApiAT10 did not result in any measurable impairment of gametocyte development or morphology, indicating the dispensability of these individual *Pf*ApiATs for the sexual stage development of the parasite (Figure 6, Figure S5).

**Figure 4:**
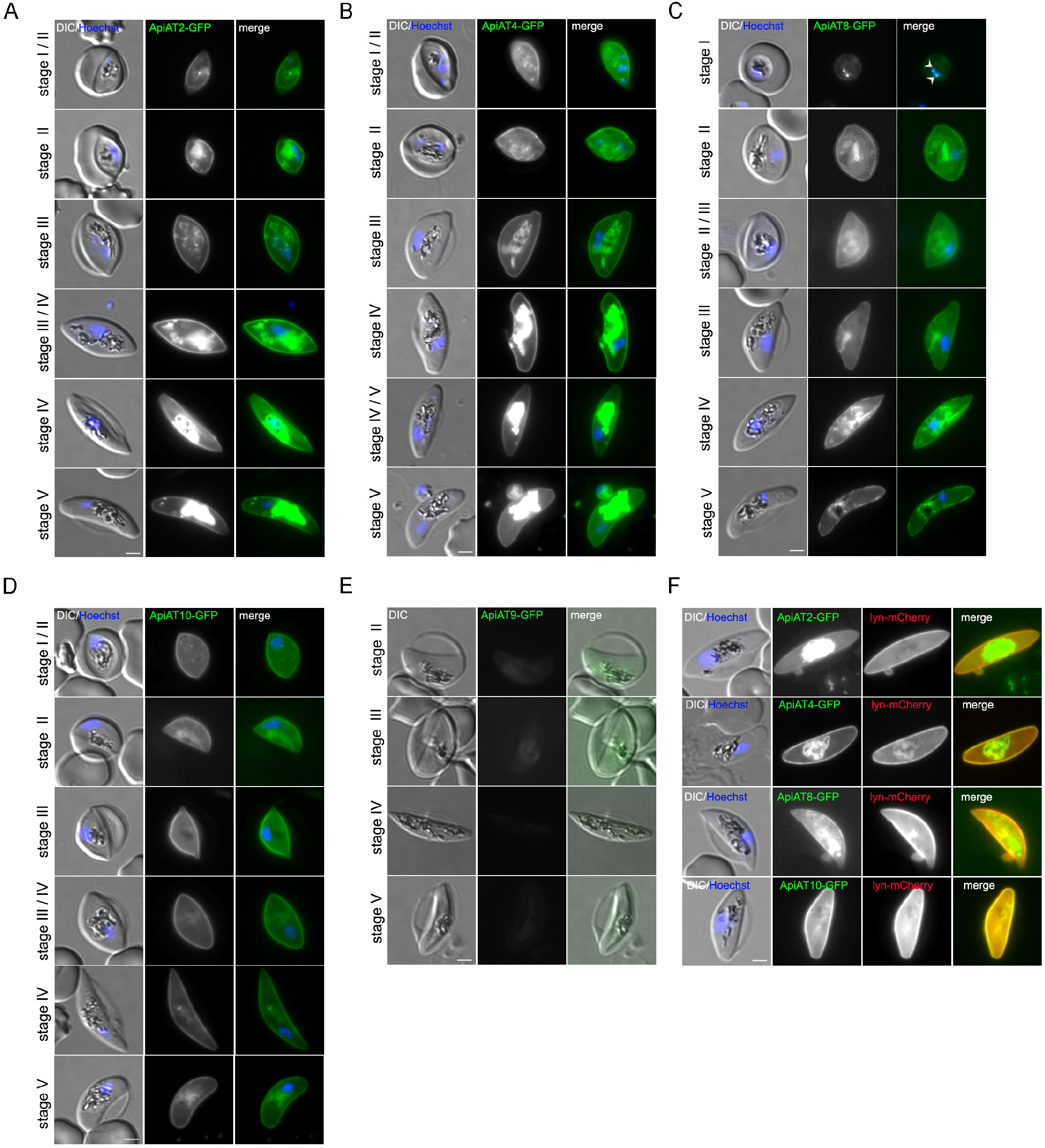
*P. falciparum* ApiATs localize to the parasite plasma membrane (PPM) during gametocyte development. **(A–D)** Localization of **(A)** *Pf*ApiAT2-GFP, **(B)** *Pf*ApiAT4-GFP, **(C)** *Pf*ApiAT8-GFP and **(D)** *Pf*ApiAT10-GFP in individual 3D7-iGP-ApiAT-GFP cell lines during gametocyte development (stage I - V) as determined by live-cell microscopy. White arrow heads indicate remaining GDV1-GFP signal observed in close proximity to the Hoechst signal in the 3D7-iGP-ApiAT8-GFP cell line **(C)**, as previously described^59,82,83^. **(E)** Localization of *Pf*ApiAT9-GFP during gametocytogenesis was assessed with the 3D7-ApiAT9-GFP cell line (see Figure 1) upon induction with choline depletion. **(F)** Co-localization of the GFP-tagged ApiAT fusion proteins with the PPM marker protein lyn-mCherry. Nuclei were stained with Hoechst-33342. Scale bar, 2 µm.

**Figure 5:**
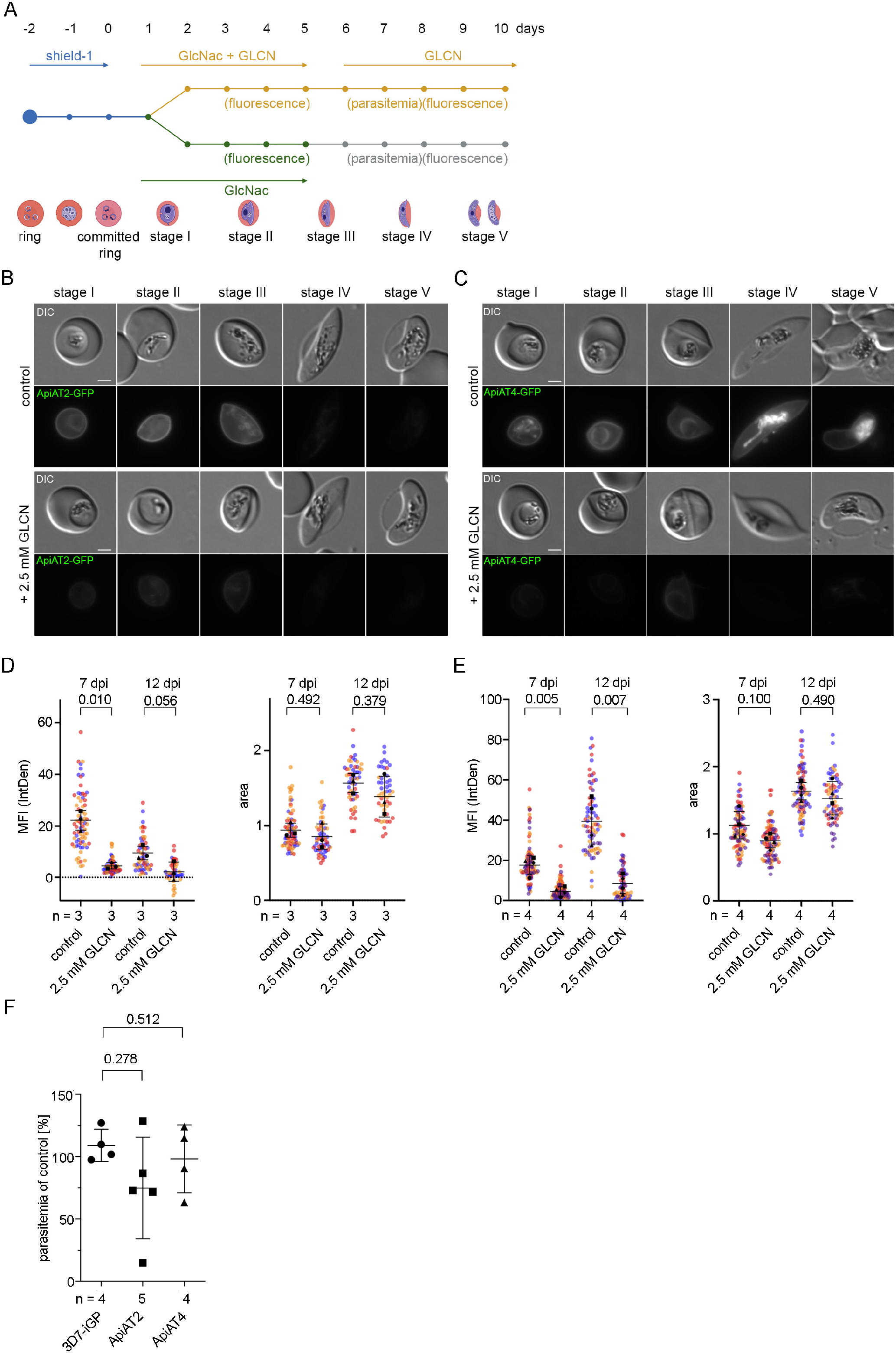
Conditional knockdown of *Pf*ApiAT2 and *Pf*ApiAT4 reveals dispensability for gametocyte development. **(A)** Schematic representation of the experimental setup. **(B, C)** Live-cell microscopy of parasites with identical settings of **(B)** 3D7-iGP-ApiAT2-GFP and **(C)** 3D7-iGP-ApiAT4-GFP stage I – V gametocytes. Scale bar, 2 µm. **(D, E)** Quantification of knockdown by measuring mean fluorescence intensity (MFI) density and size (area) of **(D)** 3D7-iGP-ApiAT2-GFP **(E)** 3D7-iGP-ApiAT4-GFP parasites at day 7 and day 12 post induction of gametocytogenesis cultured either with or without (control) 2.5 mM GLCN. Scale bar, 2 µm. Data are displayed as mean +/- SD of three (3D7-iGP-ApiAT2-GFP) or four (3D7-iGP-ApiAT4-GFP) independent experiments and individual data points are displayed on a scatterplot color-coded by experiments according to Superplots guidelines^81^. P-values displayed were determined with a two-tailed unpaired t-test with Welch’s correction. **(F)** For each condition gametocytemia at day 10 post gametocyte induction was determined by counting between 702-7693 (mean 2210) cells per condition in Giemsa-stained thin blood smears. The relative gametocytemia values (%) displayed were obtained by dividing the gametocytemia of glucosamine-treated cultures by the gametocytemia of the corresponding untreated cultures. Displayed are means +/- SD of independent growth experiments with the number of experiments (n) indicated. A two-tailed unpaired t-test with Welch’s and Benjamini-Hochberg correction was used to calculate multiplicity adjusted p-values for ApiAT2-GFP-glmS or ApiAT4-GFP-glmS versus 3D7-iGP parasites all cultured with 2.5mM GLCN.

**Figure 6:**
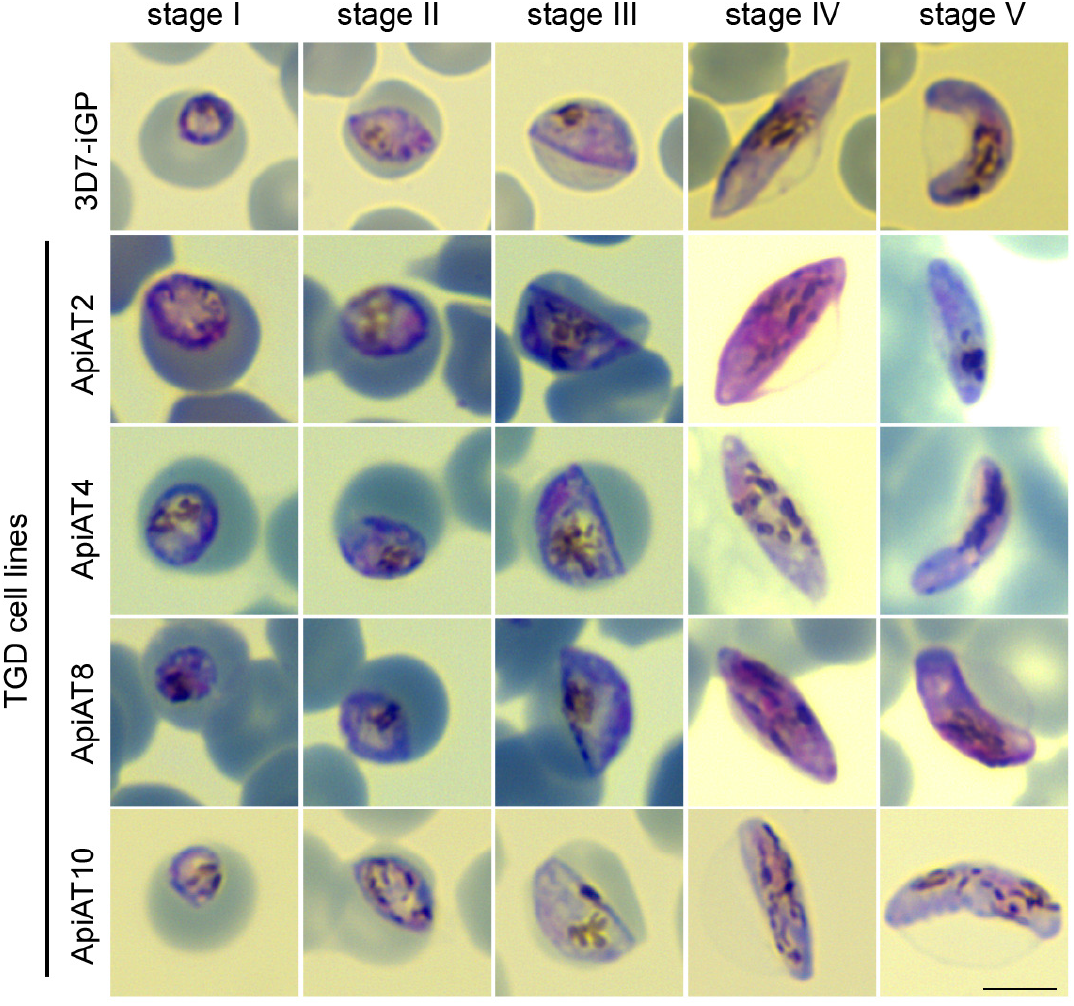
Targeted gene disruption (TGD) of *Pf*ApiAT2, *Pf*ApiAT4, *Pf*ApiAT8 and *Pf*ApiAT10 reveals dispensability of ApiATs for gametocyte development. Representative images from two (*Pf*ApiAT2, *Pf*ApiAT4) or three (*Pf*ApiAT8, *Pf*ApiAT10) independent experiments derived from Giemsa-stained thin blood smears of gametocyte stages I-V of 3D7-iGP, 3D7-iGP-ApiAT2-TGD, 3D7-iGP-ApiAT4-TGD, 3D7-iGP-ApiAT8-TGD, 3D7-iGP-ApiAT10-TGD parasites. Scale bar, 5 µm.

## DISCUSSION

We localized four putative amino acid transporters (*Pf*ApiAT2, *Pf*ApiAT4 *Pf*ApiAT8, P*f*ApiAT10) of the ApiAT-family to the PPM in asexual blood stage parasites and gametocytes. Due to the low expression of *Pf*ApiAT9-GFP – in agreement with the transcript levels of *apiat9* in these stages^37,42,43^ – no conclusive localization could be delivered. The observed PPM localization of the investigated ApiATs is in concordance with published data, since i) the *P. berghei* ApiAT8 homologue was shown to be a general cationic amino acid transporter of the PPM^29,30^, ii) *Pf*ApiAT8 has recently been localized to the PPM using an overexpression approach^31^, and iii) in *T. gondii* several ApiATs (*Tg*ApiAT1, *Tg*ApiAT2, *Tg*ApiAT3-1, *Tg*ApiAT3-2, *Tg*ApiAT3-3, *Tg*ApiAT5-3, *Tg*ApiAT6-1 and *Tg*ApiAT6-3) have been located at the PPM as well^25,29,30,32,33,44^. Recent work using episomally overexpressed *Pf*ApiAT10-GFP implied an association of this transporter with the mitochondrial membrane^36^ in *Pf*Dd2 parasites. This localization differs from the observed PPM localization of endogenously GFP-tagged *Pf*ApiAT10 in both, 3D7 as well as 3D7-iGP parasites (Figure 1B, 4C, S2A) reported in this study. We confirmed the PPM localization of *Pf*ApiAT10 by its overexpression either as a GFP or mCherry fusion using two different promoters (*crt*^45^, *ama1*^46^). In line with that, the reported reduced sensitivity to the mitochondrial electron transport chain inhibitor Atovaquone^47^ upon knockout of *Pf*ApiAT10^36^, was not observed upon targeted gene disruption in our study (Figure S2B). It is possible but appears unlikely that the reported mitochondrial association as well as reduced sensitivity to atovaquone upon overexpression is due to the different parasite strains (*Pf*Dd2^48^ versus 3D7 this study), given that *PfDd2_apiat10* (PFDd2_030017500) has only one silent mutation at position G1080A compared to *Pf3D7_apiat10*^49^.

During the intraerythrocytic development of the parasite, the amino acid requirements are largely covered by degradation of the globin polypetide^14,50^, although – for instance – the import of isoleucine is crucial for the survival of the parasite as *P. falciparum* lacks the canonical pathways for its biosynthesis^51^ and adult human hemoglobin lacks this amino acid. Dedicated amino acid transporter could fill this gap. Therefore, we tested the impact of functional inactivation of individual *Pf*ApiATs on parasite growth in asexual and sexual blood stages of *P. falciparum*. We only observed a minor but significant reduction in parasite growth upon knockdown or gene disruption of *Pf*ApiAT2 and *Pf*ApiAT4 in asexual blood stages without compensatory up-regulation of other *Pf*ApiATs on the transcriptional level, as indicated by qPCR analysis. The phenotypes are in agreement with a previously reported reduced parasite multiplication rate of 36 % in *Pb*ApiAT4 knockout parasites^35^. Moreover our data is also in line with the finding that *Pb*ApiAT8 is not essential for asexual replication^29,30,35,52^. This might be explained by functional redundancy either within the ApiAT family, as recently observed for *T. gondii*^34^, or by the presence of yet unassigned transporters capable of transporting essential amino acids such as isoleucine across the PPM. Like in *T. gondii*, overlapping substrate specificities and lower transport levels might be sufficient for parasite growth *in vitro*^34^. Of note, in *T. gondii* the arginine transporter *Tg*ApiAT1 has been shown to be differently regulated on the translational level in dependence of arginine mediated by an upstream open reading frame (uORF) present on the 5’ leader sequence of the transcript^26^. A similar layer of regulation might also be present in *Pf*ApiATs. However, since *Toxoplasma* and *Plasmodium* only share one ApiAT, the likely most ancestral ApiAT2^25^, the regulatory elements as well as the general characteristics and substrates of the different *Pf*ApiATs remain largely unknown.

Interestingly, a knockout of the ApiAT8 homologue of *P. berghei* (PBANKA_0208300) resulted in strongly reduced number of mature gametocytes with an aberrant morphology of the remaining parasites^30^ and strongly reduced exflagellation^35^. In our study, functional inactivation of *Pf*ApiAT8 via targeted gene disruption had no impact on gametocyte development and morphology, which might reflect the pronounced differences in gametocyte development between the rodent infecting *P. berghei* and the human infecting *P. falciparum* parasites^53^. For future work, it will be interesting to target multiple ApiATs by gene disruption in parallel to assess their putative synergy and to probe into *Pf*ApiAT function in other fast-replicating stages of *P. falciparum* such as liver stages, for which the essentiality of several metabolic processes has recently been shown and that primarily rely on the amino acid uptake from their host^54^.

## METHODS

### *P. falciparum* culture

Blood stages of *P. falciparum* 3D7^55^ were cultured in human red blood cells (O+ or B+). Cultures were maintained at 37°C in an atmosphere of 1 % O_2_, 5 % CO_2_ and 94 % N_2_ using RPMI complete medium containing 0.5 % Albumax according to standard protocols^56^. In order to obtain highly synchronous parasite cultures late schizonts were isolated by percoll gradient^57^ and cultured with fresh erythrocytes for 4 hours. Afterwards sorbitol synchronization^58^ was applied in order to remove remaining schizonts resulting in a highly synchronous ring stage parasite culture with a four-hour age window.

Induction of gametocytogenesis was done as previously described^41,59^. Briefly, GDV1-GFP-DD expression was achieved by addition of 2 or 4 µM Shield-1 to the culture medium and gametocyte cultures were treated with 50 mM N-acetyl-D-glucosamine (GlcNAc) for five days starting 72 h post Shield-1 addition to eliminate asexual parasites^60^. Alternatively, asexual ring stage cultures with >10 % parasitemia, cultured in the presence of choline, were synchronized with Sorbitol^58^ and washed twice in choline-free RPMI medium. Cells were kept in choline-free medium for the entirety of the assay. After one reinvasion cycle cultures at trophozoite stage were treated with 50 mM N-acetyl-D-glucosamine (GlcNAc)^60^ and kept on this for five days. Gametocytes were maintained in RPMI complete medium containing 0.25 % Albumax and 0.25 % sterile filtered human serum (Interstate Blood Bank, Inc. Memphis, TN, USA).

Growth assays in low amino acid medium were performed using amino acid restricted RPMI medium prepared as previously described^20^. Briefly, complete medium was added in a 1/20 dilution to amino acid-free RPMI medium 1640 (US Biological, R9010-01). This resulted in a 1:20 of the concentration of every amino acid compared to the standard RPMI complete based medium.

### Cloning of plasmid constructs for parasite transfection

For endogenous tagging using the SLI system^38^ and glmS based conditional knockdown^40^ a 855 (*Pf*ApiAT4/MFR5/PF3D7_1129900) and 1001 bp (*Pf*ApiAT2/MFR4/PF3D7_0914700) homology region was amplified using 3D7 gDNA and cloned into pSLI-PIC1-GFP-glmS^61^ using the NotI/MluI restriction site.

For endogenous GFP-tagging a 886 bp (*Pf*ApiAT8/NPT1/PF3D7_0104800), 953 bp (*Pf*ApiAT9/MFR2/PF3D7_0104700), 881 bp (*Pf*ApiAT10/MFR3/PF3D7_0312500) homology region was amplified using 3D7 gDNA and cloned into pSLI-GFP^38^ using the NotI/MluI restriction site.

For SLI-based targeted gene disruption (SLI-TGD) a 396 bp (*Pf*ApiAT4, *Pf*ApiAT8), 402 bp (*Pf*ApiAT9), 393 bp (*Pf*ApiAT10) and 435 bp (*Pf*ApiAT2) homology region was amplified using 3D7 gDNA and cloned into the pSLI-TGD plasmid^38^ using NotI and MluI restriction sites.

For overexpression constructs the full length *Pf*ApiAT10 sequence was amplified from parasite gDNA and cloned into pARL-ama1^46^-AIP-mCherry-yDHODH^61^ using the XhoI/KpnI restriction site or into the pARL-crt^45^-PF3D7_0324600-GFP-hDHFR^37^ plasmid using the KpnI/AvrII restriction site. Oligonucleotides used to generate the DNA fragments are summarized in Table S1.

For co-localization experiments the plasmid pLyn-FRB-mCherry^38^ was used.

### Western blot analysis

Immunoblots were performed using saponin-lysed, infected erythrocytes. Parasite proteins were separated on a 12 % SDS-PAGE gel as described previously^62,63^ and transferred to a nitrocellulose membrane (Amersham Protran; 0.45-µm pore nitrocellulose membrane; GE Healthcare) using a Trans-Blot device (Bio-Rad) according to the manufacturer’s instructions. The membranes were blocked with 3 % skim milk in TBS for 30 minutes and then probed with mouse anti-GFP (clone 7.1 and 13.1; 1:1,000, Roche) or rabbit anti-aldolase^64^ (1:2,000). The chemiluminescent signal of the horseradish peroxidase-coupled secondary antibodies (Dianova) was visualized using a Chemi Doc XRS imaging system (Bio-Rad) and processed with Image Lab 5.2 software (Bio-Rad).

To perform loading controls and ensure equal loading of parasite material, rabbit anti-aldolase^64^ antibodies were used. The corresponding immunoblots were incubated twice in stripping buffer (0.2 M glycine, 50 mM dithiothreitol, 0.05 % Tween 20) at 55°C for 1 h and washed 3 times with Tris-buffered saline for 10 min before re-probing.

### Transfection of *P. falciparum*

For transfection, Percoll-purified^57^ parasites at late schizont stage were transfected with 50 µg plasmid DNA using Amaxa Nucleofector 2b (Lonza, Switzerland) as previously described^65^. Transfectants were selected using either 4 nM WR99210 (Jacobus Pharmaceuticals), 0.9 µM DSM1^66^ (BEI Resources), or 2 μg/mL blasticidin S (Life Technologies, USA). In order to select for parasites carrying the genomic modification via the SLI system^38^, G418 (ThermoFisher, USA) at a final concentration of 400 µg/mL was added to a culture with about 5 % parasitemia. The selection process and integration test were performed as previously described^38^.

### Imaging

All fluorescence images were captured using a Zeiss Axioskop 2plus microscope with a Hamamatsu Digital camera (Model C4742-95) or a Leica D6B fluorescence microscope equipped with a Leica DFC9000 GT camera and a Leica Plan Apochromat 100x/1.4 oil objective.

Microscopy of live parasite-infected erythrocytes was performed as previously described^67^. Briefly, parasites were incubated in standard culture medium with 1 µg/mL Hoechst-33342 (Invitrogen) for 15 minutes at 37°C prior to imaging. 5.4 µL of infected erythrocytes were added on a glass slide and covered with a cover slip. Nuclei were stained with 1 µg/mL Hoechst-33342 (Invitrogen). Mitochondria were visualized by incubation of parasites with 20 nM MitoTracker Red 665 CMXRos (Invitrogen) for 15 min at 37°C prior to imaging. Images were processed using Fiji^68^ and Adobe Photoshop CC 2021 was used for display purposes only.

### Growth assay

For growth assays of TGD cell lines a flow cytometry assay, adapted from previously published assays^69,70^, was performed to measure proliferation over five days. For growth under low amino acid conditions TGD and wild type cell lines were cultured in parallel in standard and amino acid-depleted medium for five days. Each day parasite cultures were resuspended and 20 µL samples were transferred to an Eppendorf tube. 80 µL RPMI containing Hoechst-33342 and dihydroethidium (DHE) was added to obtain final concentrations of 5 µg/mL and 4.5 µg/mL, respectively. Samples were incubated for 20 min (protected from UV light) at room temperature, and parasitemia was determined using an LSRII flow cytometer by counting 100,000 events using the FACSDiva software (BD Biosciences) or using an ACEA NovoCyte flow cytometer.

### Gametocyte quantification assay

Giemsa-stained blood smears at day 10 post induction of GDV1 expression were obtained and at least 10 fields of view were recorded using a 63x objective per treatment and time point. Erythrocyte numbers were then determined using the automated Parasitemia software (http://www.gburri.org/parasitemia/) while the number of gametocytes was determined manually in >700 erythrocytes per sample.

### GlmS-based knockdown

GlmS-based knockdown assay was adapted from previously published assays^40,71^. To induce knockdown, highly synchronous early ring stage parasites were split in two dishes, 2.5 mM glucosamine was added to one of them and parasite growth was measured by FACS after two and four parasite replication cycles. Parasite cultures were inspected daily by Giemsa smears and, if necessary, diluted to avoid growth bias caused by high parasitemia. As an additional control, the same amount of glucosamine was also added to 3D7 wildtype parasites. For all analyses, medium was changed daily, and fresh glucosamine was added every day. GlmS-based knockdown assay was adapted from previously published assays^40,71^. To induce knockdown, highly synchronous early ring stage parasites were split into two dishes, 2.5 mM glucosamine was added to one of them and parasite growth was measured by FACS after two and four parasite replication cycles. Parasite cultures were inspected daily by Giemsa smears and, if necessary, diluted to avoid growth bias caused by high parasitemia. As an additional control, the same amount of glucosamine was also added to 3D7 wildtype parasites. For all analyses, medium was changed daily, and fresh glucosamine was added every day.

Knockdown was quantified by fluorescence live cell microscopy using schizonts about 40 h post glucosamine treatment. Parasites with similar size were imaged, and fluorescence was captured with the same acquisition settings to obtain comparable measurements of the fluorescence intensity. Fluorescence intensity (integrated density) was measured with Fiji^68^, and background was subtracted in each image. The data was visualized with GraphPad Prism version 8 (GraphPad Software, USA).

For knockdown experiments in gametocytes synchronized ring stage cultures were induced by the addition of Shield-1, as described above. At day 3 post induction the culture was spilt into two dishes and one dish was cultured in the presence of 2.5 mM glucosamine for the remaining ten days. Knockdown was quantified by fluorescence live cell microscopy at day 7 and 10 post induction, as described above.

### Drug assays

Drug assays were adapted from previously described assays^72–74^. Briefly, 3D7-iGP and 3D7-iGP-ApiAT10-TGD parasites were synchronized to a 4 h time window resulting in 0–4 h ring stage parasites. At 24 hpi, parasitemia was determined by flow cytometry and the drug susceptibility assays were set up in black 96-well microtiter plates (Thermo Scientific) with 0.1 % starting parasitemia and 2 % hematocrit in a final volume of 200 µl. In each plate, infected erythrocytes in the absence of drugs treated with DMSO only served as positive controls, while uninfected RBCs served as negative controls (for background subtraction). Parasites were incubated with varying concentrations of dihydroartemisinin (DHA) (Adipogen, Switzerland, AG-CN2-0468) (0–50 nM) and Atovaquone (Cayman, Item No.23802) (0–16 nM).

After 96 h of incubation, parasite growth was determined by measuring the fluorescence of SYBR Gold (Invitrogen). Therefore, 100 µl/well supernatant was discarded without disturbing the RBC layer and 100 µl of lysis buffer (20 mM Tris, 0.008 % saponin, 0.08 % Triton X-100, 1x SYBR Gold) was added to each well. Plates were incubated in the dark for 2 h at room temperature before measuring fluorescence using the EnVision Multimode Plate Reader (PerkinElmer), as described previously^74^. In order to calculate IC_50_ values, the measured values were normalized to the uninfected erythrocytes and plotted in GraphPad Prism version 8 (GraphPad Software, USA) as % of DMSO control. Dose-response curves were generated using nonlinear regression (curve fit > dose-response inhibition > (log) inhibitor vs. normalized response—variable slope).

### Quantitative real-time PCR (qPCR)

Parasites at different time-points (8, 16, 32 and 44 hpi) were harvested for 3D7-ApiAT2-TGD, 3D7-ApiAT4-TGD, 3D7-ApiAT8-TGD, 3D7-ApiAT10-TGD and 3D7-WT to obtain RNA samples for quantitative real-time PCR. Highly synchronous ring stage parasite cultures were grown for another 40 h and TRIzol samples were harvested in the following cycle. Volumes of prewarmed TRIzol used for infected erythrocyte lysis and storage of RNA samples depended on the parasite stage: ring stages were lysed in 5x volumes, trophozoites in 10x volumes and schizonts in 20x volumes of the settled cell pellet. RNA was purified and checked for absence of genomic DNA. cDNA synthesis with random hexamers and quantitative real-time PCR was performed exactly as previously described^75^. Primers for each of the *apiat* genes, for genes to control for parasite stages (*sbp1*^76^, *tom22, ama1*) and for housekeeping genes (*arginyl-tRNA synthetase*^75^, *fructose bisphosphate aldolase*^77^) are listed in Table S1. Amplification efficiencies of the primer pairs were determined over six log10 dilutions of gDNA (10 ng – 0.0001 ng) and were shown to have similar values between 1.915 and 2.001 (Table S1). Expression of *apiat* genes and controls were analyzed in relation to expression of the *arginyl-tRNA synthetase* gene (normalizer).

### Software

Schematic protein representations were designed using IBS^78^, predicted protein domains were obtained from plasmodDB^79^ inferred from TMHMM^80^. Parasite icons were generated using BioRender (biorender.com). Statistical analyses were performed with GraphPad Prism version 8 (GraphPad Software, USA).

## Acknowledgements

We thank Till Voss for 3D7-iGP parasites, Jacobus Pharmaceuticals for WR99210 and Greg Burri for the parasitemia software. DSM1 (MRA-1161) was obtained from MR4/BEI Resources, NIAID, NIH.

## Author contribution

Conceptualization: JSW, JS, TWG, AB; Methodology: JSW, EP, PCB, PMR, TS, JS, TWG, AB; Investigation: JSW, CVG, GF, JMR, EP, JLF, HVT, SS, PCB; Formal Analysis: JS, AB; Writing original manuscript: JSW, AB; Review & Editing: JSW, TWG, AB; Funding Acquisition: TWG, AB, JS; Resources: PMR, TS; Project Administration: TWG, AB; Supervision: TWG, AB. All authors read and approved the manuscript.

## Figures

**Figure S1: PCR analysis and western blots of 3D7-ApiAT-GFP and 3D7-ApiAT-TGD cell lines. (A)** PCR-based analysis of unmodified wildtype (WT) and transgenic knock-in (KI) cell lines (3D7-ApiAT2-GFP-glmS, 3D7-ApiAT4-GFP-glmS, 3D7-ApiAT8-GFP, 3D7-ApiAT9-GFP, 3D7-ApiAT10-GFP). Specific genomic modifications resulting from correct integration of the respective SLI-based vectors were tested targeting the 3’- and 5’-end of the locus. **(B)** Western blot analysis of wildtype (3D7) and knock-in (KI) cell lines (3D7-ApiAT2-GFP-glmS, 3D7-ApiAT4-GFP-glmS, 3D7-ApiAT8-GFP, 3D7-ApiAT10-GFP) using mouse anti-GFP to detect the tagged full-length protein (upper panel) and rabbit anti-aldolase to control for equal loading (lower panel). Protein sizes are indicated in kDa. **(C)** PCR-based analysis of unmodified wildtype (WT) and transgenic parasites modified by targeted gene disruption (TGD).

**Figure S2: *Pf*ApiAT10 is PPM localized and its functional inactivation has no** influence on Atovaquone sensitivity. (A) Live-cell microscopy of 3D7-iGP-ApiAT10-GFP parasites stained with MitoTracker Red CMXRos across the IDC and in gametocytes. Stages as indicated: R = ring stage, ET = early trophozoite, LT = late trophozoite, ES = early schizont, LS = late schizont, G = gametocyte; Nuclei were stained with Hoechst-33342. Scale bar, 2 µm. **(B)** Drug susceptibility assays of 3D7-iGP and 3D7-iGP-ApiAT10-TGD parasites were performed with Atovaquone (left) and dihydroartemisinin (DHA, right). Parasite growth was determined by measuring the DNA content using SYBR gold when exposed to varying concentrations of drugs for 96 h. The growth of DMSO-treated control parasites was set to 100%. Shown are means +/- SD of 4 or 5 independent biological replicates performed in technical duplicates. Calculated IC_50_ values with 95% confidence intervals are shown above each graph.

**Figure S3: Quantitative real-time PCR of individual *Pf*ApiAT-encoding genes and controls in 3D7-ApiAT-TGD and wildtype parasites**. Expression of *apiat* genes and control genes (*sbp1, tom22, ama1, fructose-bisphosphate aldolase*) normalized to the housekeeping control *arginyl-tRNA synthetase* over four different time points during intraerythrocytic development cycle of *P. falciparum* parasites. Bars with individual measurements from n=2 (3D7-ApiAT2-TGD, 3D7-ApiAT4-TGD) or n=3 (3D7-ApiAT8, 3D7-ApiAT10, 3D7 wildtype) biological replicates. Gene accession numbers, primer sequences and amplification efficiencies are listed in Table S1.

**Figure S4: PCR analysis of 3D7-iGP-ApiAT-GFP cell lines and Giemsa smears from *Pf*ApiAT2 and *Pf*ApiAT4 conditional knockdown experiments. (A)** PCR-based analysis of unmodified wildtype (WT) and transgenic knock-in (KI) cell lines (3D7-iGP-ApiAT2-GFP-glmS, 3D7-iGP-ApiAT4-GFP-glmS, 3D7-iGP-ApiAT8-GFP, 3D7-iGP-ApiAT10-GFP). **(B)** Giemsa smears of stage I – V 3D7-iGP-ApiAT2-GFP-glmS and 3D7-iGP-ApiAT4-GFP-glmS gametocytes cultured either without (control) or with 2.5 mM glucosamine (GLCN). Scale bar, 5 µm. **(C)** Live cell imaging of 3D7-iGP stage IV and V gametocytes. Nuclei were stained with Hoechst-33342. Scale bar, 2 µm. **(D)** Live cell microscopy of 3D7-iGP-ApiAT2 stage IV and V gametocytes from Figure 5B, with adjusted brightness. Scale bar, 2 µm.

**Figure S5: PCR analysis of 3D7-iGP-ApiAT-TGD cell lines and Giemsa smears from 3D7-iGP-ApiAT8-TGD stage IV and V gametocytes showing normal morphology. (A)** PCR-based analysis of unmodified wildtype (WT) and transgenic TGD cell lines (3D7-iGP-ApiAT2-TGD, 3D7-iGP-ApiAT4-TGD, 3D7-iGP-ApiAT8-TGD, 3D7-iGP-ApiAT10-TGD). (**B**) Giemsa smears of 3D7-iGP-ApiAT8-TGD and 3D7-iGP stage IV and V gametocytes from three independent experiments. Scale bar, 5 µm.

**Table S1: Oligonucleotides used for cloning and quantitative real-time PCR (qPCR)**.

