## Supplementary figures and images for "Characterization of apicomplexan amino acid transporters (ApiATs) in the malaria parasite *Plasmodium falciparum*"

### Figure S1

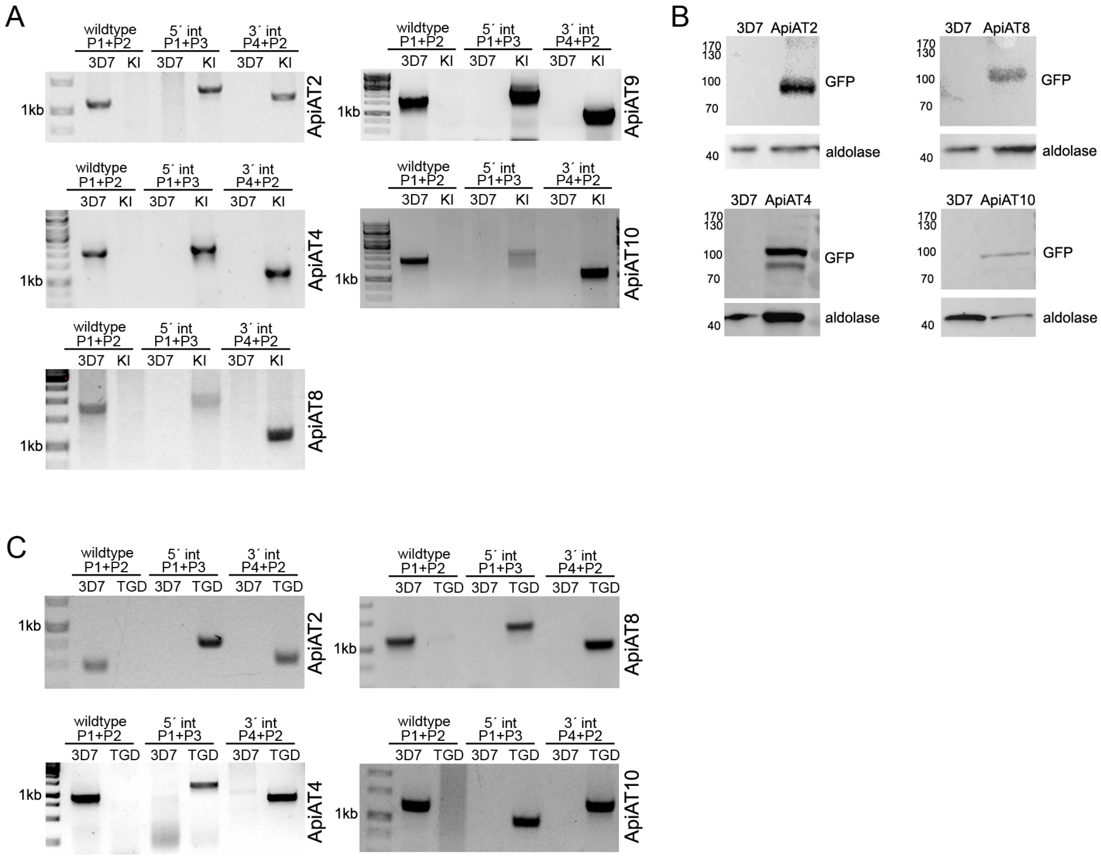

### Figure S2

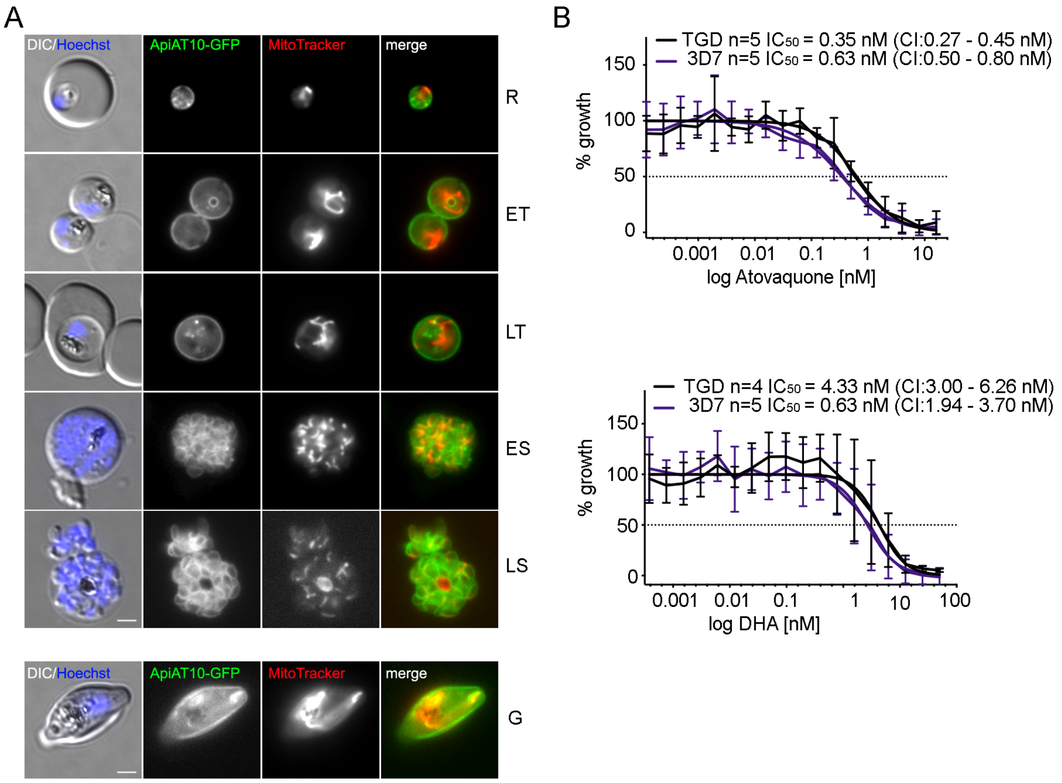

### Figure S3

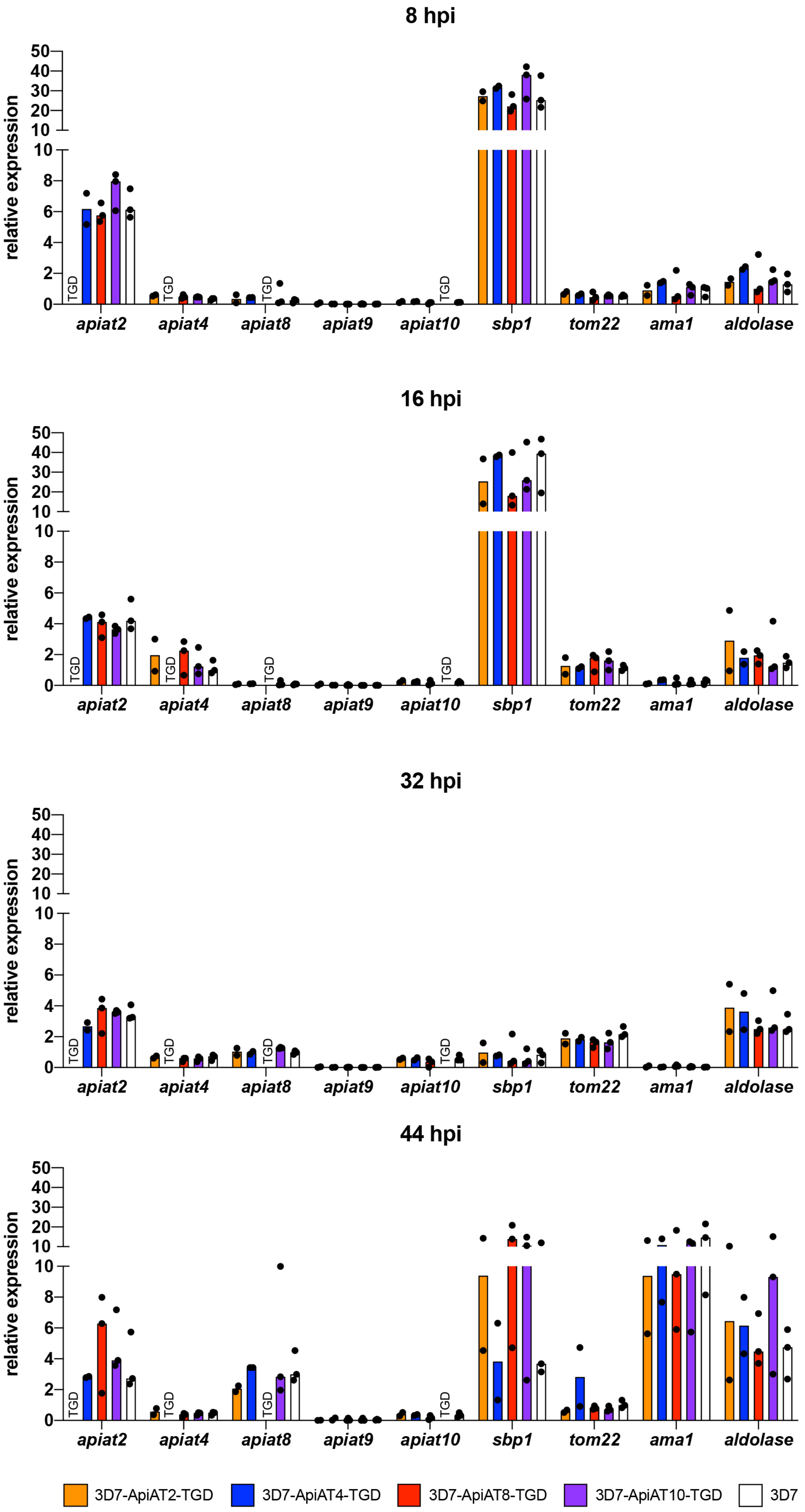

### Figure S4

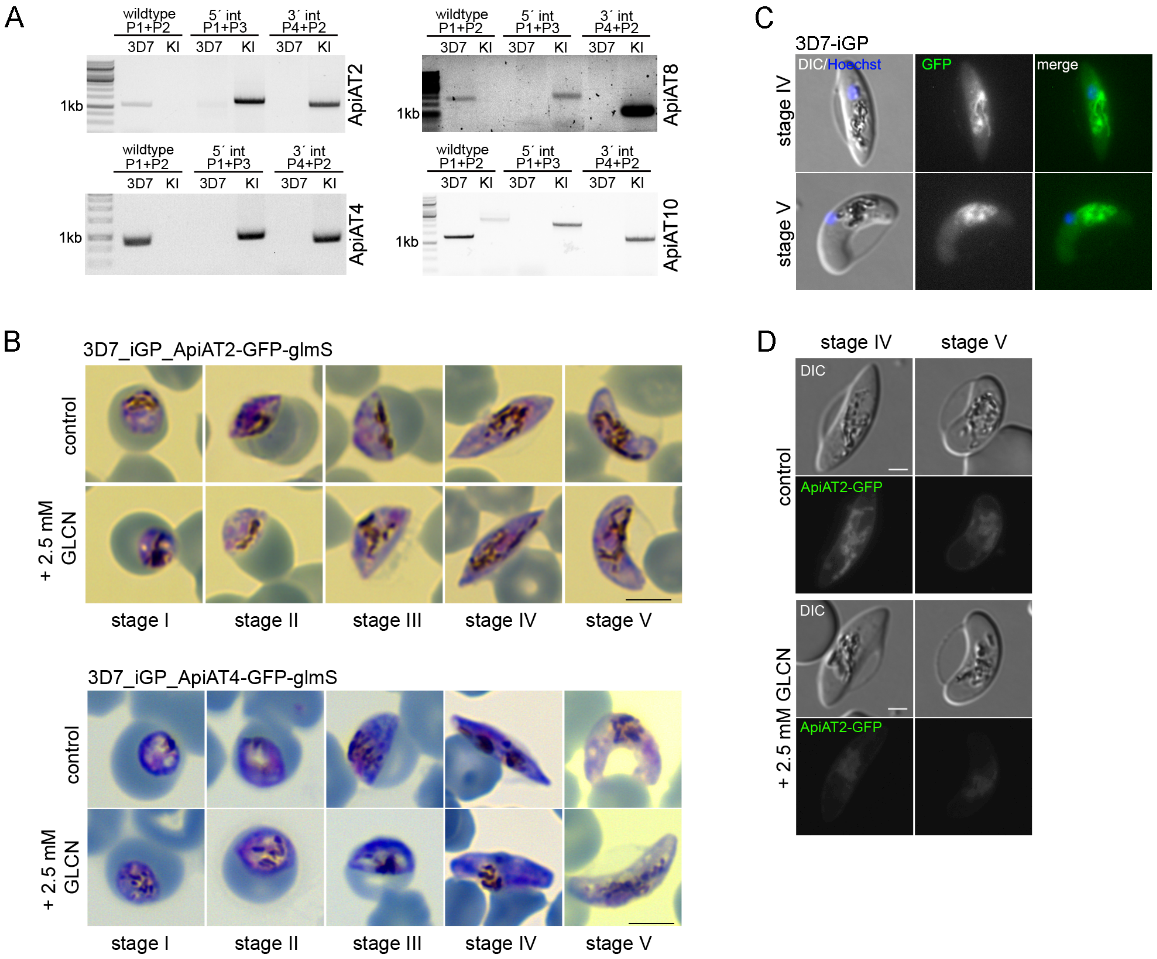

### Figure S5

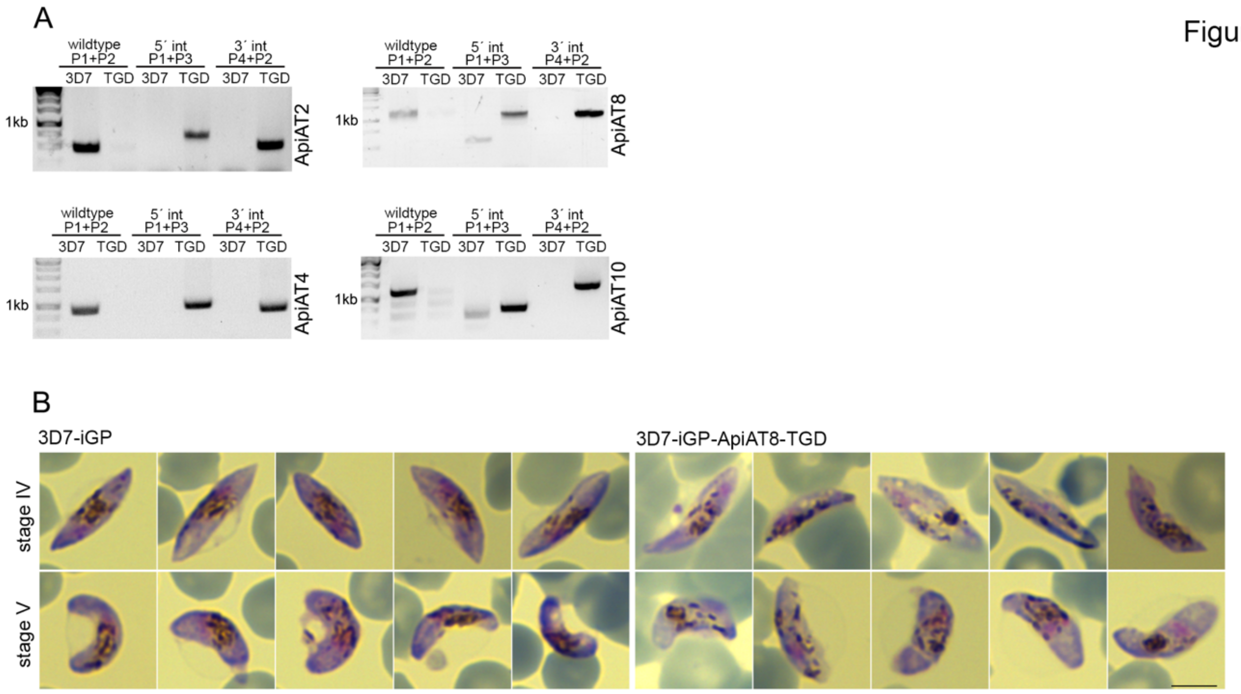
