## Supplementary material for "Characterization of apicomplexan amino acid transporters (ApiATs) in the malaria parasite *Plasmodium falciparum*": Table S1

**Table S1: Oligonucleotides used for cloning and quantitative real-time PCR (qPCR).**

| ID | Target or Primer name | Sequence | Efficiency | Purpose |
|---|---|---|---|---|
| 286 | HR ApiAT4 fw | GGGacgcgtCTTTGATGATGCGTCGTCTCCATAG | | cloning pSLI-GFP |
| 287 | HR ApiAT4 rv | GGGgcggccgctaaGACCATGATAAGAAAAAAGATGGAC | | |
| 329 | HR ApiAT8 fw | GGGgcggccgctaaGGTAGATAAAGATAGTACAGA | | |
| 330 | HR ApiAT8 rv | GGGacgcgtATTCATTGGGAGTTTATCCTTTTCTCC | | |
| 331 | HR ApiAT9 fw | GGGgcggccgctaaGCCAAATTATATATACTAATAGCTCAATG | | |
| 332 | HR ApiAT9 rv | GGGacgcgtCTTTTTGGTTTGACATCTTTTGTAT | | |
| 333 | HR ApiAT10 fw | GGGgcggccgctaaGTGTGGATGATATAAACAATTCATC | | |
| 334 | HR ApiAT10 rv | GGGacgcgtCTTTGAAGAAGGAAGGGAAATCATGTGAT | | |
| 454 | HR ApiAT2 fw | GGGgcggccgctaaGTTCAGCGAGATCTACTGGATTTGC | | |
| 455 | HR ApiAT2 rv | GGGacgcgtGTTTTTGTTTTTCTTCTTTTCTGTAAC | | |
| 321 | HR ApiAT4-TGD fw | GCGGCCGCTAAAGTTCCAAAAAAAAGTCTACAATTATATTG | | cloning pSLI-TGD |
| 322 | HR ApiAT4-TGD rv | acgcgtATCATATTCTCCATGCTCATTTTTTTGACA | | |
| 466 | HR ApiAT2-TGD fw | GGGgcggccgctaaGCGTCAGATGTATCAAAGGAAAATAT | | |
| 467 | HR ApiAT2-TGD rv | GGGacgcgtAAAGGCTAAAGCTAAAATAATTTGTC | | |
| 340 | HR ApiAT8-TGD fw | gcggccgctaaAGTAACTCAATATTCCATAAG | | |
| 341 | HR ApiAT8-TGD rv | acgcgtCCAACAAATAATATTAAATAACTGCCC | | |
| 344 | HR ApiAT10-TGD fw | gcggccgctaaGTAAAGGACAAAACAATAATCTCCC | | |
| 345 | HR ApiAT10-TGD rv | acgcgtGACTAATGATATTTTATTCGTT | | |
| 303 | intcheck ApiAT4 fw | CGAAAAGGGAGATGAGTTCAAATGATAATG | | integration check PCR |
| 304 | intcheck ApiAT4 rv | TATATATATATATAACCCTTTCGTTACTGG | | |
| 359 | intcheck ApiAT9 fw | GGATACTTATAGGATTACAAAAGGG | | |
| 360 | intcheck ApiAT9 rv | GACGTGTTCCCTTGAACTAATCCACG | | |
| 361 | intcheck ApiAT8 fw | AGTAACTCAATATTCCATAAG | | |
| 362 | intcheck ApiAT8 rv | GTATGTATGTATAATTGTTCGATATGG | | |
| 363 | intcheck ApiAT10 fw | GTAAAGGACAAAACAATAATCTCCC | | |
| 364 | intcheck ApiAT10 rv | CATATATAAAGAGGTACGTAAGAATAG | | |
| 483 | intcheck ApiAT2 fw | CCAGGACAAGAATCCTTAATTTTTGG | | |
| 484 | intcheck ApiAT2 rv | CCTGTTCATATGATCGTACACGTCAG | | |
| 373 | intcheck ApiAT4-TGD fw | CTTCAGTTGTGTTTATTTTTGGTAT | | |
| 374 | intcheck ApiAT4-TGD rv | CATTTCCTGGAAATAATCCTGATGC | | |
| 379 | intcheck ApiAT8-TGD fw | GTTTACATTAAAAATTGTACTCTGTATG | | |
| 380 | intcheck ApiAT8-TGD rv | CTATCTTTATCTACCATTCCGCTTGC | | |
| 383 | intcheck ApiAT10-TGD fw | CCTTGTTAATAATAAACATGAAAAA | | |
| 384 | intcheck ApiAT10-TGD rv | CCCCAAATACATTATTTTCATATATG | | |
| 485 | intcheck ApiAT2-TGD fw | ACTTCATTTTATTCCTTTTTG | | |
| 486 | intcheck ApiAT2-TGD rv | CCATACGTATGAGAATTTTAAAATGGC | | |
| 226 | GFP rv | TTTTGTTGATAATGGTCTGC | | |
| | GFP as 272 | CCTTCGGGCATGGCACTC | | |
| 238 | pARL sense 55 | GGAATTGTGAGCGGATAACAATTTCACACAGG | | |
| 456 | 456_ApiAT2_for | GTCAAGCACCCAACGACCTA | 1.949 | qPCR |
| 457 | 457_ApiAT2_rev | TGCCATCCACTAAATCCACCA | | |
| 458 | 458_ApiAT9_for | GGGATGAAAAATCTTTCCAATTCGT | 1.961 | |
| 459 | 459_ApiAT9_rev | AGTTTATGTGCCTCCATGGTAA | | |
| 460 | 460_ApiAT4_for | TCAGCATGTGCAAATGGACAATTA | 2.001 | |
| 461 | 461_ApiAT4_rev | CCACACCACACACTAGGTTCA | | |
| 462 | 462_ApiAT10_for | AGGCAATAGCTCAAGGGCTC | 1.962 | |
| 463 | 463_ApiAT10-rev | TGAAGAAGGAAGGGAAATCATGTG | | |
| 464 | 464_ApiAT8_for | ACCAGGTGCCAAACAAAAGAC | 1.982 | |
| 465 | 465_ApiAT8_rev | TATATGCACGCTGAGGTCGC | | |
| | arginyl-tRNA synthetase_for (75) | TTCAAAACACGAAGTGGAACAAC | 1.915 | |
| | arginyl-tRNA synthetase_rev (75) | AATTCTCTGCAGCAAGTCGC | | |
| | fructose-bisphosphate aldolase_for (77) | TGTACCACCAGCCTTACCAG | 1.938 | |
| | fructose-bisphosphate aldolase_rev (77) | TTCCTTGCCATGTGTTCAAT | | |
| | sbp1_for (76) | TTAGCCGACGAACCAACACA | 1.916 | |
| | sbp1_rev (76) | TTCGGTTGTCTCTGGTACTGCA | | |
| | tom22_for | GCCCATAAGGATGCCATTCG | 1.961 | |
| | tom22_rev | CACCTGCTATCCATACAACCCA | | |
| | ama1_for | TGGGTAATCCATGGACGGAA | 1.967 | |
| | ama1_rev | TGAGTTCCAGCTACTTCAGCA | | |
| JSW 75 | ApiAT10 crt fw | GGGggtaccATGAAAAAAGTAAAGGACAAAAC | | cloning pARL-*crt* ApiAt10-GFP |
| JSW 76 | ApiAT10 crt rv | GGGcctaggCTTTGAAGAAGGAAGGGAAATCATG | | |
| JSW 53 | ApiAT10 ov fw | GGGctcgagATGAAAAAAGTAAAGGACAAAAC | | cloning pARL-*ama1* ApiAt10-GFP |
| JSW 54 | ApiAT10 ov rv | GGGggtaccCTTTGAAGAAGGAAGGGAAATCATG | | |
